# A simple, cost-effective microfluidic device using a 3D cross-flow T-junction for producing decellularized extracellular matrix-derived microcarriers

**DOI:** 10.1101/2024.10.30.621138

**Authors:** Farah Kamar, Connor J. Gillis, Grace Bischoff, Anorin Ali, John R. de Bruyn, Lauren E. Flynn, Tamie L. Poepping

## Abstract

Cell therapies using human mesenchymal stromal cells (MSCs) are promising for a wide variety of clinical applications. However, broad-scale clinical translation is limited by conventional culture methods for MSC expansion within 2D tissue-culture flasks. MSC expansion on ECM-derived microcarriers within stirred bioreactor systems offers a promising approach to support MSC growth. Previously, our team established methods for fabricating ECM-derived microcarriers from a variety of decellularized tissue sources using electrospraying techniques. However, these microcarriers are relatively large and have a broad size distribution, which may limit their utility. Smaller and more uniform microcarriers may be favorable for MSC growth within bioreactors and have greater potential to serve as a minimally invasive injectable cell delivery platform. To address these limitations, the current project focused on the development of a new microfluidic-based approach enabling both uniform and small microcarrier production. Using a novel, modified 3D T-junction design, we successfully generated microcarriers using human decellularized adipose tissue (DAT) as the ECM source. Our new cost-effective device produced microbeads that were small and monodisperse, at a range of flow rate combinations and with high production rates. Photo-crosslinking using rose bengal allowed for the generation of microcarriers that were stable following rehydration, with a mean diameter of 196 ± 47 µm. Following methods optimization and microcarrier characterization, *in vitro* studies confirmed that the new microcarriers supported human adipose-derived stromal cell (hASC) attachment and growth, as well as ECM production, across 14 days within spinner flask bioreactors. Overall, this study demonstrates the feasibility of using our novel, cost-effective, and reusable microfluidics device to generate cell-supportive microcarriers comprised exclusively of ECM in a size range that could be injected via a small gauge needle and were stable in long-term culture.

## 1 INTRODUCTION

Cell therapies using human mesenchymal stromal cells (MSCs) have shown promise for a wide range of clinical applications. However, a limitation for broad-scale clinical translation is that conventional culture methods for MSC expansion within 2D tissue-culture flasks can lead to cell senescence, reflected through changes in cell shape, diminished proliferative and differentiation capacities, and an altered secretome [1]. New approaches are critically needed to enable MSC expansion while preserving, or ideally augmenting, their pro-regenerative functionality. Notably, there is evidence that dynamic culture within bioreactor systems may be favorable for directing the lineage-specific differentiation of MSCs [2], [3], [4], as well as enhancing their secretion of pro-regenerative paracrine factors [5], [6]. In addition, providing MSCs with a substrate that more closely resembles the biochemical and biophysical properties of the extracellular matrix (ECM) within their native niche may be favorable for counteracting cell senescence [7].

Combining these strategies, MSC expansion on ECM-derived microcarriers within stirred bioreactor systems offers a promising approach to support MSC growth [8]. Previously, our team has established methods for fabricating microcarriers comprised exclusively of ECM using a variety of decellularized tissue sources, including decellularized adipose tissue [4], [9], cartilage [10], dermis [8], myocardium [8], and brain [11]. In brief, the microcarriers were generated by electrospraying an ECM suspension into liquid nitrogen to form spherical beads which remained stable without chemical crosslinking following lyophilization and controlled rehydration [8], [12]. The microcarriers generated with this approach are typically quite large, however, with mean diameters ranging from ∼450 μm [12] to over 900 μm [11], depending on the ECM source, and it can be challenging to control the size distribution. While few studies have systemically compared the effects of microcarrier size, it is recognized that the microcarrier diameter can influence MSC attachment and growth [13], [14], [15], [16], with the majority of commercially available microcarriers being in the size range of 100 – 300 μm [17]. An added benefit to microcarriers in this size range is that they have the potential to serve as a minimally invasive injectable cell delivery platform via small gauge needles.

Recognizing the limitations of our previous technology, the primary focus of this study was on establishing a new cost-effective method for fabricating ECM-derived microcarriers sourced from decellularized tissues, which had a mean diameter of ∼200 μm and were more uniformly sized. Microfluidics offers a potential avenue for smaller, monodisperse microcarrier production [18]. In particular, T-junction microfluidic devices can create monodispersed droplets (<1-3% dispersity) [19] using two intersecting channels of immiscible fluids – composed of a dispersed phase (DP) and a continuous phase (CP) – which allows for the creation of microcarriers through cross-flow shearing forces. The size and production rate of the microcarriers are easily altered by changing the channel flow rates [20]. Previously, a 3D T-junction microfluidic device was fabricated by Yoon *et al*. to produce collagen type I beads that were < 200 µm in diameter [21]. Their device was fabricated with a CP channel (500 x 500 μm^2^ in cross-section) intersecting a smaller DP channel (50 x 50 μm^2^ in cross-section) and hence described as a 3D T-junction. While the overall design concept of the 3D T-junction is promising for yielding small microcarriers, this type of device is prone to clogging and the fabrication process is complicated and costly as it requires nanofabrication facilities and micron-level alignment precision.

In the current study, we used a micro-milling method for microfluidic device fabrication. Our CP channel was micro-milled into an acrylic material and a 34-gauge needle was directly inserted into the CP channel to create the intersecting DP channel. This fabrication method is much less complicated, and the device is readily reusable, making it far more cost-effective than previous methods. The new microfluidic device design allows for easier fabrication and bead production through a modified 3D T-junction design. An important design feature is that the needle tip can be inserted into the midplane of the continuous-phase crossflow to increase the drag and shearing forces experienced by the dispersed phase to optimize bead production size and dispersity. As proof-of-concept, we generated microcarriers using human decellularized adipose tissue (DAT) as the ECM source; DAT is readily available as surgical waste and has been demonstrated to have cell-supportive and pro-regenerative properties [22], [23], [24]. Following methods optimization and microcarrier characterization, *in vitro* studies were performed to confirm that the new microcarriers supported MSC attachment and expansion within spinner flask bioreactors. These tests were performed using human adipose-derived stromal cells (hASCs) as the matching tissue-specific MSC source for the DAT.

## 2 METHODS

### 2.1 Materials

Unless otherwise stated, all chemicals and reagents were purchased from Sigma Aldrich Canada and were used as received.

### 2.2 Microfluidic device fabrication

A 500 x 500 μm^2^ channel measuring 40 mm in length was micro-milled into a 12-mm thick rectangular block of acrylic, creating an open-bottomed CP channel. At the 15-mm mark along the channel, a hole with inner diameter of 190 μm was drilled through the top of the acrylic block into the center of the 500 μm-wide channel, followed by a shallow counterbore for the tip of the needle hub. A blunt-tipped 34G needle with an inner diameter of 80 μm was inserted into this small hole, creating the intersecting DP channel. The 34G needle was precisely inserted midway into the CP channel, such that DP would be injected into the mid-plane of the flow as depicted in Fig. 1. Barbed tubing micro-connecters were inserted into the ends of the CP channel and tubing was secured to create inlets and outlets. The bottom of the CP channel was sealed by screwing a 110-mm diameter acrylic plate (compatible with a microscope stage opening) to the top chamber with 16 screws placed around the channel. The screws were countersunk into the plate to avoid abrasion with any surfaces, especially microscope lenses used for analysis. This design allowed the device to be reopened for cleaning if the channels clogged. In addition, the DP channel needle can be easily replaced, making the device reusable. This new fabrication method also avoids potential alignment issues that could arise in designs requiring the alignment of separate top and bottom sections to form a 3D microchannel. Figure 1 shows the final device.

**Fig. 1.**
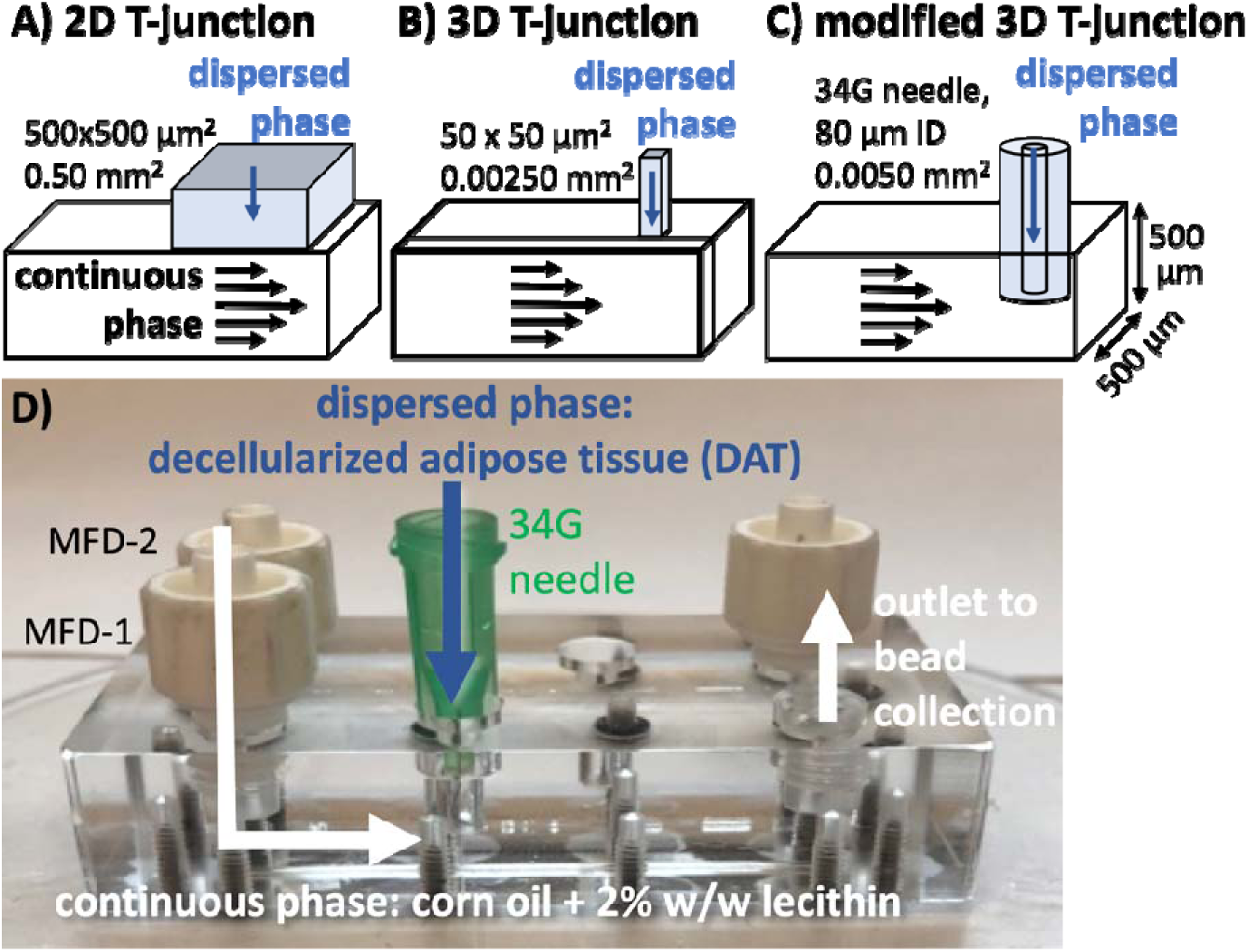
A modified 3D T-junction using a micro-milled continuous-phase channel and 34G needle as dispersed-phase channel serves as a low-cost, reusable microbead generator with enhanced shearing forces. (A-C) A comparison of microfluidic device T-junction designs: (A) 2D T-junction, (B) 3D T-junction used by Yoon *et al.* [21], formed by joining two moulded layers, and (C) modified 3D T-junction with midstream injection as designed and used for this study. Arrows portray the fluid flow directions of the two fluids including the horizontal continuous phase (CP) in black and the vertical dispersed phase (DP) in blue. (D) A photograph of the microfluidic device used in this study where the CP consisted of corn oil with 2% w/w lecithin as surfactant, and the DP consisted of the decellularized adipose tissue (DAT) suspension. The shallow bore for the needle hub was precisely milled to facilitate inserting the needle end precisely into the CP channel midplane. The device shown has two parallel microbead generators in the same device for efficiency. Cross-sectional dimensions of the DP channel are given for comparison.

### 2.3 Tissue procurement and decellularization

Discarded human adipose tissue was collected with informed consent from scheduled plastic and reconstructive surgeries provided through surgical collaborators in London, ON, with human research ethics board approval from Western University (HSREB # 105426). Fresh tissue was transported on ice in sterile phosphate buffered saline (PBS, Wisent Bioproducts) with 0.3 mM bovine serum albumin (BSA, Bio-shop) and decellularized following a published detergent-free process [25]. The DAT was then lyophilized for 48 h and stored until further use. DAT from a minimum of 5 donors was pooled and processed to generate a DAT suspension through published protocols [4] involving cryomilling, enzymatic digestion with 1% w/w α-amylase, and homogenization in 0.2 M acetic acid at a concentration of 50 mg/mL (based on the initial dry mass). The resulting stock DAT suspension was stored at 4°C for up to 1 month prior to further use.

### 2.4 DAT viscosity

The viscosity of the DAT suspension was measured using a strain-controlled rotational shear rheometer (MCR302, Anton Paar) with a concentric cylinder (CC27) tool. The viscosity was measured at DAT concentrations of 10 mg/mL, 20 mg/mL, and 30 mg/mL, with the stock diluted in 0.2 M acetic acid (Fisher Scientific). For each concentration, 20 mL of DAT suspension was transferred into the rheometer cup. Data were recorded (RheoPlus, Anton Paar) for shear rates ranging from 0.1 s^-1^ to 100 s^-1^, at two points per decade, and acquired for 800 s at the lowest shear rate and 10 s at the highest shear rate. Viscosity was measured at 4°C, 15°C, 22°C, and 37°C for each DAT concentration, allowing up to a few minutes for the suspension to reach within 0.1°C of the desired temperature before measurements were taken. Each measurement was repeated three times.

### 2.5 Microbead production

To fabricate the microbeads, for the DP fluid, a DAT suspension at a concentration of 20 mg/mL was loaded into one syringe. For the CP fluid, a second syringe was filled with corn oil (Fisher Scientific) mixed with 2% w/w lecithin (Santa Cruz Biotechnology Inc.) as a surfactant to minimize bead agglomeration after production [21]. Both syringes were secured onto a computer-controlled, low-pressure syringe pump (neMSYS 290N with Qmix Elements software, CETONI GmbH). Initial testing revealed that straining was required to prevent the DAT suspension from clogging and allow it to flow smoothly through the microfluidic device needle tip. To this end, the suspensions were filtered consecutively through cell strainers of 100-μm and 70-μm mesh size. The DAT and corn-oil syringes were attached to the DP channel and CP channel, respectively, of our T-junction device using 3-mm diameter plastic tubing. The DAT suspension was maintained at ∼22^°^C for easier flow.

To view the microbeads being formed in the microchannels, the microfluidic device was positioned on the stage of an inverted microscope (Olympus IX71) and illuminated using a broad-spectrum light source (X-Cite 120 metal halide lamp). A 2-mm length of the CP channel was viewed using a 20X objective lens, and images were acquired using a CMOS camera (Phantom VEO 340L, Vision Research). ImageJ software was used to analyze the diameter of the microbeads. To analyze the relationship between microbead diameter and flow rate, flow rates of 50, 100, and 250 μL/min were used in the DP channel (Q_DP_), while flow rates of 200, 500, and 1500 μL/min were used in the CP channel (Q_CP_). The coefficient of variation (standard deviation/mean) was used to assess bead size variability.

Further processing (i.e., crosslinking, rehydration, cell culturing) and analyses were performed on beads made with flow rates of Q_DP_=100 μL/min and Q_CP_=500 μL/min. The DAT microbeads in corn oil were collected in a beaker surrounded by dry ice, to induce freezing to avoid coalescence and loss of bead shape. Once the batch was completed, 50 mL of rose bengal in acetone (0.25% w/v) was added to the beaker as a photo-crosslinking agent [9], [26], [27] A salt-water ice bath was prepared at a temperature between -5°C and -10°C. In this temperature range, the DAT beads remained frozen and retained their shape during crosslinking while the oil remained liquid, allowing the rose bengal to penetrate the beads. The microbead beaker was moved from the dry ice to this saltwater bath and illuminated for 6 hours with a broadband white light source to allow for chemical crosslinking of the collagen within the microbeads [9]. The crosslinked microbeads were washed with 100% ethanol in a 100-μm pore cell strainer and then transferred to a 50 mL conical tube and slowly rehydrated using an ethanol series diluted in PBS (99%, 98%, 95%, 90%, 80%, 70%, 60%, 50%, 25%, 12.5%, 0%) over the span of 2 days. After rehydration, the microbeads were tested in terms of their performance as microcarriers.

### 2.6 Microcarrier size distribution

Microcarriers were imaged using an EVOS XL Core Imaging System. The diameter was computed using ImageJ to plot the size distribution and determine the average microcarrier diameter (n = 100 microcarriers per batch, N = 2 independent batches).

### 2.7 Scanning electron microscopy

Microcarrier surface ultrastructure was assessed using scanning electron microscopy (SEM). To prepare the samples for imaging, rehydrated microcarriers were incubated for 1 h in a 50 mL conical tube at room temperature (∼22°C) in a solution of 2.5% SEM-grade glutaraldehyde in PBS under agitation at 100 RPM on an orbital shaker, followed by static incubation for 24 h at 4□. Next, the microcarriers were rinsed 3 times with deionized water and lyophilized. Prior to imaging, the samples were mounted on stubs, coated with osmium, and imaged using a LEO 1530 SEM at an accelerating voltage of 2 kV and working distance of 6 mm.

### 2.8 Mechanical testing

To determine the Young’s modulus of the microcarriers, compression testing was performed using a CellScale MicroTester system equipped with a 154-μm diameter cantilever within a PBS bath at 37□. Microcarriers (n = 6 microcarriers/batch, N = 3 independent batches) were compressed to 50% of their original diameter at a strain rate of 0.01 s^-1^. Three cycles were used for preconditioning the microcarriers, and then data were collected over the next 3 cycles and used to determine the Young’s modulus based on published methods [28].

### 2.9 *In vitro* characterization using human adipose-derived stromal cells

Following validation that the microfluidic device reproducibly generated DAT microcarriers that were stable after photo-chemical crosslinking, proof-of-concept *in vitro* studies were performed to validate that the new microcarriers supported human MSC attachment and growth within a spinner flask bioreactor system. Applying a tissue-specific approach, hASCs were selected as the cell source to test the DAT microcarriers. hASCs were isolated using previously described and published methods [25] and cultured in DMEM/F12 media (Wisent Bioproducts) supplemented with 10% fetal bovine serum and 1% penicillin-streptomycin (Life Technologies). hASCs at passage 2-3 were used for microcarrier seeding within a CELLSpin spinner system (Integra Biosciences), following published protocols [9]. In brief, 100-mL spinner flasks were inoculated with 5.3 g of microcarriers (hydrated mass) in 50 mL of complete medium and incubated overnight at 37□ and 5% CO_2_. The next day, each flask was seeded with 2.5 x 10^6^ hASCs following an established intermittent-stirring seeding protocol [9]. In the first trial, the flasks were stirred continuously after seeding at 20 RPM for a period of 14 days. Due to microcarrier aggregation observed at 20 RPM, three additional trials were performed at 40 RPM, comparing the response of hASCs from three different donors (N=3). Half-media changes were performed every 2-3 days.

### 2.10 Characterization of cell attachment and growth

Microcarrier samples were collected at 24 h, 72 h, 7 d, and 14 d after seeding to assess cell attachment using LIVE-DEAD® staining (Life Technologies) following manufacturer’s instructions. Samples were visualized using an LSM 800 confocal microscope (Zeiss Canada) [29]. At each time point, additional samples were collected, frozen, and lyophilized for analysis of double-stranded DNA (dsDNA) content via the PicoGreen® assay as a quantitative measure of total cell content to assess cell growth over time. In brief, the lyophilized samples were digested overnight using Proteinase K in tris-EDTA buffer (TE Buffer) and the DNA was extracted using the DNeasy® Blood & Tissue Kit (Qiagen), following the manufacturer’s protocols. Samples were then analyzed with the PicoGreen® assay using a CLARIOstar® plate reader, following the manufacturer’s instructions.

### 2.11 Biochemical assays

The hydroxyproline and dimethylmethylene blue (DMMB) assays were performed as a measure of the total collagen content and sulphated glycosaminoglycan (sGAG) content of the baseline microcarriers, as well as the hASC-seeded microcarriers after 14 days of culture within the CELLSpin flasks to probe ECM remodeling. For both assays, lyophilized microcarriers were digested with Proteinase K in TE buffer for 2 hours under agitation at 300 RPM and 56°C, then incubated for 10 min at 95°C to inactivate the enzyme. The assays were then performed following published protocols [10] and using a CLARIOstar® plate reader.

### 2.12 Statistical analyses

All data are presented as the group mean ± standard deviation. All statistical analyses were performed using GraphPad Prism. Data were analyzed using unpaired t-test or two-way ANOVA with Tukey’s post-hoc test for multiple comparisons, unless otherwise noted. Differences were reported as statistically significant at p<0.05. All graphs were created using data analysis software (GraphPad Prism v. 8, La Jolla, CA; or MATLAB 2021b, MathWorks, MA, USA) as noted. Linear regression analysis was performed on the plot of microbead diameter as a function of the ratio of Q_DP_/Q_CP_ (MATLAB 2021, Mathworks Inc., USA), given that previous studies have shown relative independence with Q_DP_ and inverse relationship with Q_CP_.

## 3 RESULTS

### 3.1 DAT suspension viscosity

The viscosity of the DAT suspension as a function of shear rate, concentration, and temperature was investigated (Fig. 2) for a better understanding of the material to optimize fabrication parameters. The viscosity data were fitted to the Cross model,

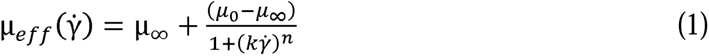

**Fig. 2.**
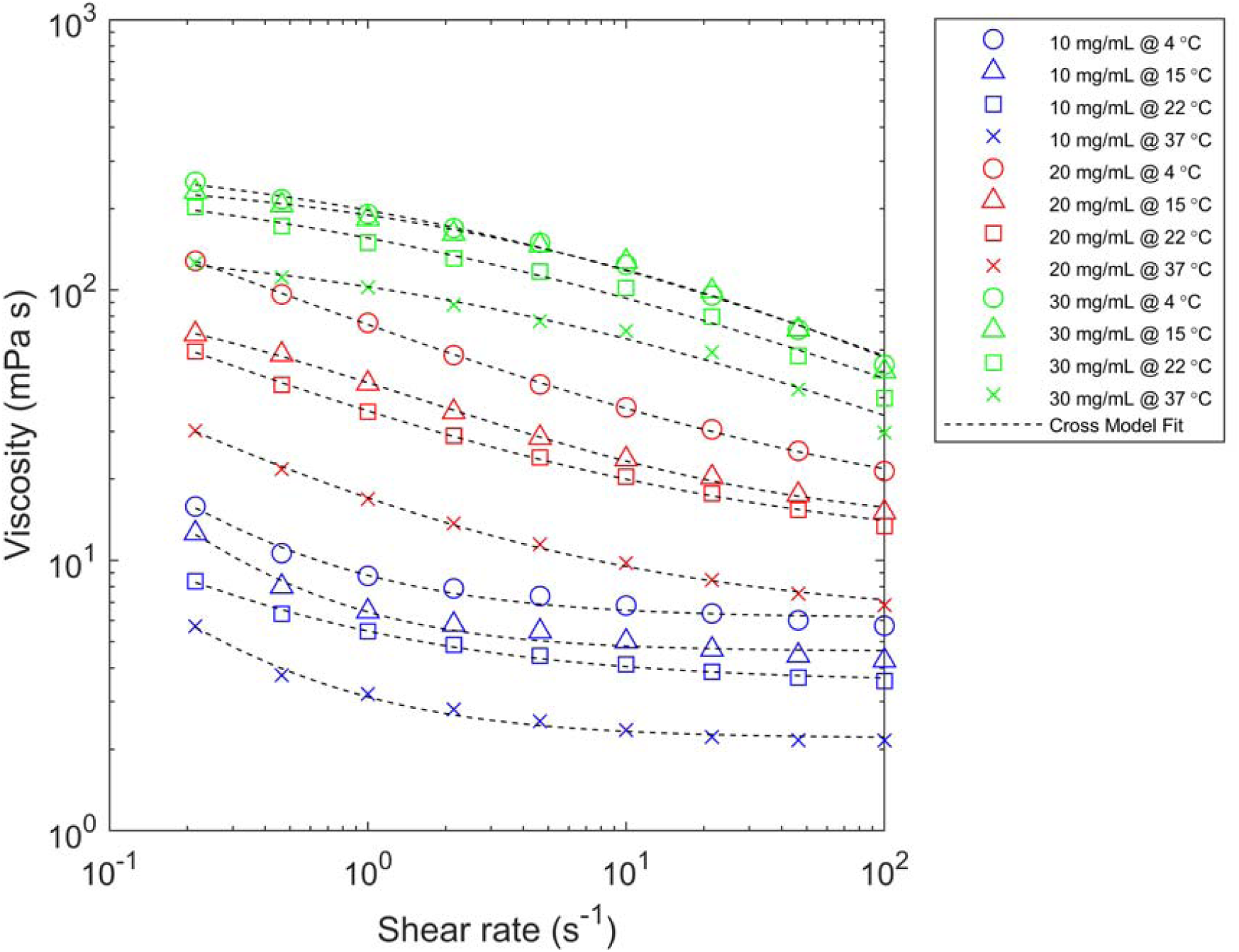
The viscosity of the DAT suspensions were well described by the Cross model as a function of shear rate for all concentrations and temperatures. Viscosity (in mPa⋅s) is shown for DAT suspension concentrations of 10 mg/mL (blue), 20 mg/mL (red), and 30 mg/mL (green) as a function of shear rate (in s^-1^), measured at 4°C, 15°C, 22°C, and 37°C for each concentration. Uncertainties based on three repeated measurements are smaller than the marker size. Dashed lines represent fits to the Cross model.

where 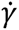 is the shear rate, (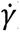), µ_oo_, and µ_O_ are the effective viscosity, infinite-shear viscosity, and zero-shear viscosity, respectively, and *k* and *n* are fluid-specific parameters. (See Supplementary Table T1 for table of fit parameters). Concentration was the strongest determinant of viscosity. At low temperatures and low shear rates, the viscosity for the 20 mg/mL suspension was ∼8-9 times higher than for 10 mg/mL, but this lessened to 3-4 times higher for high temperatures and high shear rates. The viscosity of the 30 mg/mL suspension was 20-30 times higher than the 10 mg/mL suspension at low shear rates, lessening to 10-20 times higher than 10 mg/mL for high shear rates. For each concentration, at a given temperature, the viscosity was 3-6 times higher at the lower shear rates compared to the highest rate tested, and the viscosity was 2-4 times higher at 4°C compared to 37°C. These viscosity results were then considered towards the implementation of the microbead generator, suggesting that using a high flow rate and high temperature for the dispersed phase would reduce the viscosity and facilitate working with a higher concentration of DAT as desired.

### 3.2 Microbead production

Microbead diameter and production rates are dominated by the continuous phase flow rate, Q_CP_ (Fig. 3). Microbead diameter within the microfluidic device (Fig. 3A) depended inversely on Q_CP_ (i.e., higher rates leading to smaller microcarrier sizes), with less dependence on Q_DP_, thus leading to a linear relationship with the flow ratio Q_DP_/Q_CP_ (Fig. 3B), where microbead diameters smaller than 200 µm were achieved with flow-rate ratios <0.2. Although using a high Q_CP_ of 1500 µL/min resulted in bead sizes smaller than 200 µm regardless of which Q_DP_ was used, and the highest production rates of hundreds of beads per second, it also resulted in a bimodal size distribution. All the flow rate combinations showed spherical beads. The image in Fig. 3D shows beads with a bimodal distribution having a dominant size (Bead A) and a secondary size (Bead B), while beads with a more monomodal distribution are shown in Fig. 3C. The bimodal bead size distribution approaches an equal proportion of dominant and secondary beads as Q_DP_ increases. (Fig. 3F). The secondary beads were small (<100 µm) and showed high variability in diameter (Fig. 3G), while the primary beads were more consistent in size. The variability in primary bead size at each flow rate was 5% or less when Q_CP_ was 200 µL/min or 500 µL/min and Q_DP_

**Fig. 3.**
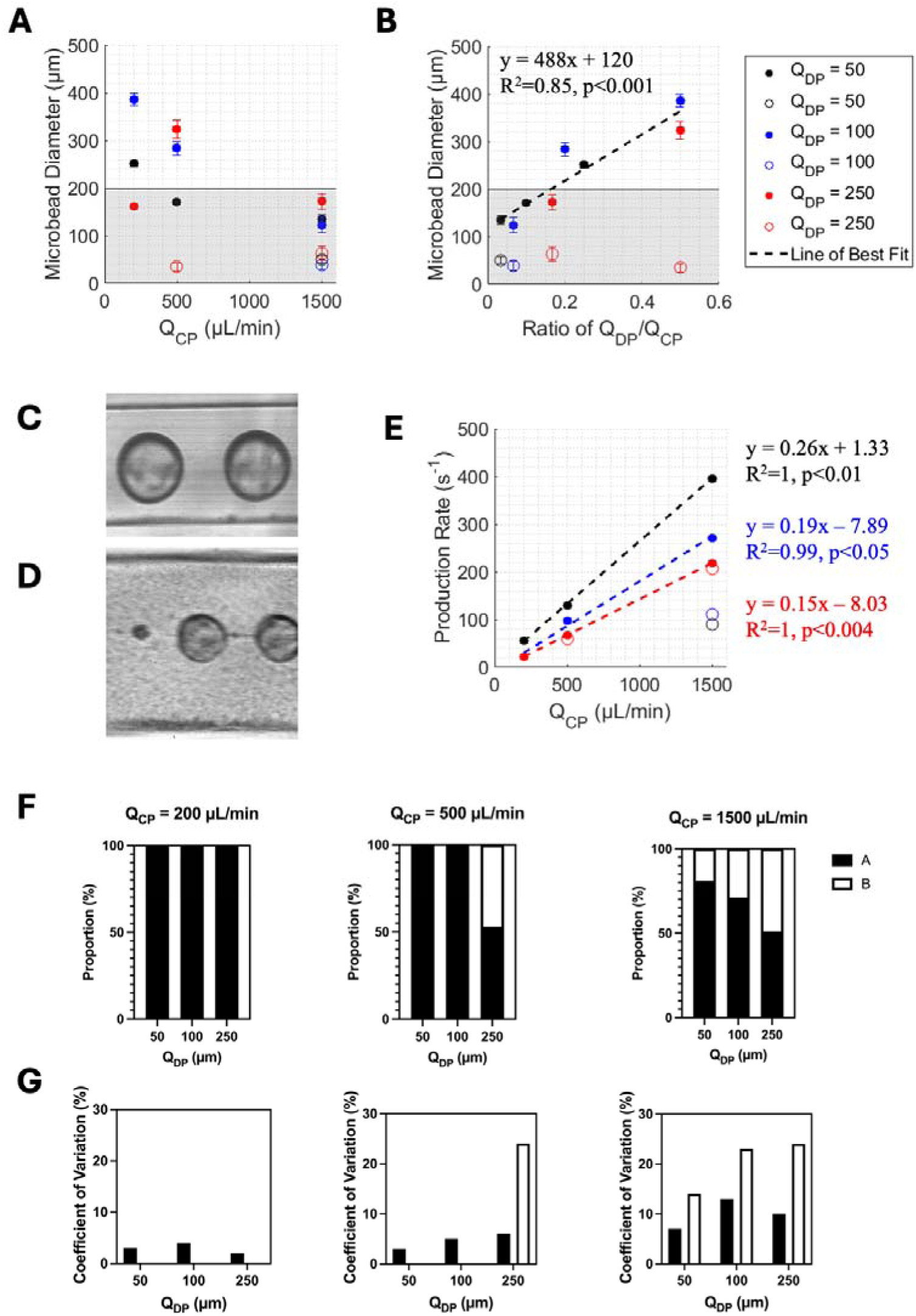
Microbead diameter and production rates across nine different flow rate combinations. (A) Microbead diameter (µm) as a function of continuous-phase flow rates (Q_CP_) using a DAT concentration of 20 mg/mL and dispersed phase flow rates (Q_DP_) of 50 µL/min (black), 100 µL/min (blue), and 250 µL/min (red). Error bars represent the standard deviation (n = 50). For flow rates displaying a bimodal distribution, the filled circle represents the dominant size (Bead A) and the open circle represents the less dominant size (Bead B). The shaded grey region indicates the flow rate combinations producing beads smaller than 200 µm. (B) Microbead diameter as a function of the ratio of Q_DP_/Q_CP_ with a line of best fit for all flow-rate ratios <1 and for dominant beads only. An example image of the (C) monomodal (Q_DP_ = 50 μL/min, Q_CP_ = 200 μL/min) and (D) bimodal (Q_DP_ = 50 μL/min, Q_CP_ = 1500 μL/min) microbead production. (E) Production rate (beads/second) is shown as a function of Q_CP_ for the three different Q_DP_ used with the lines of best fit shown for dominant beads only. (F) Proportion of Bead A and Bead B and (G) coefficient of variation (%) is shown across all flow rate combinations.

≦100 µL/min. Beads produced using Q_DP_/Q_CP_ of 100/500 were selected for further processing and study as they were monodisperse with low variability (5%), had moderately high production rates (∼100 beads/sec), and had a mean diameter of 200 µm.

### 3.3 Microcarrier physical characterization

With the selected parameters, the microcarriers (i.e. microbead after rehydration) were observed to have a relatively uniform spherical shape following rehydration (Fig. 4A). The microcarrier diameter (Fig. 4A) was determined to be 196 ± 47 µm, reproducible across batches. The microcarriers were smaller than the original microbeads before rehydration (283 ± 14 μm). Additionally, SEM imaging revealed that the microcarriers had a relatively smooth surface ultrastructure compared to our previous DAT microcarriers generated through electrospraying [4], with some regions showing a more porous network of interconnected fibers and sheet-like structures (Fig. 4B). Mechanical testing showed that the microcarriers were soft and compliant, with a linear elastic response with a Young’s modulus of 0.23 ± 0.05 kPa (N=3 batches, n=6 microcarriers/batch), which is lower than the 3–4 kPa range reported for human adipose tissue [25].

**Figure 4.**
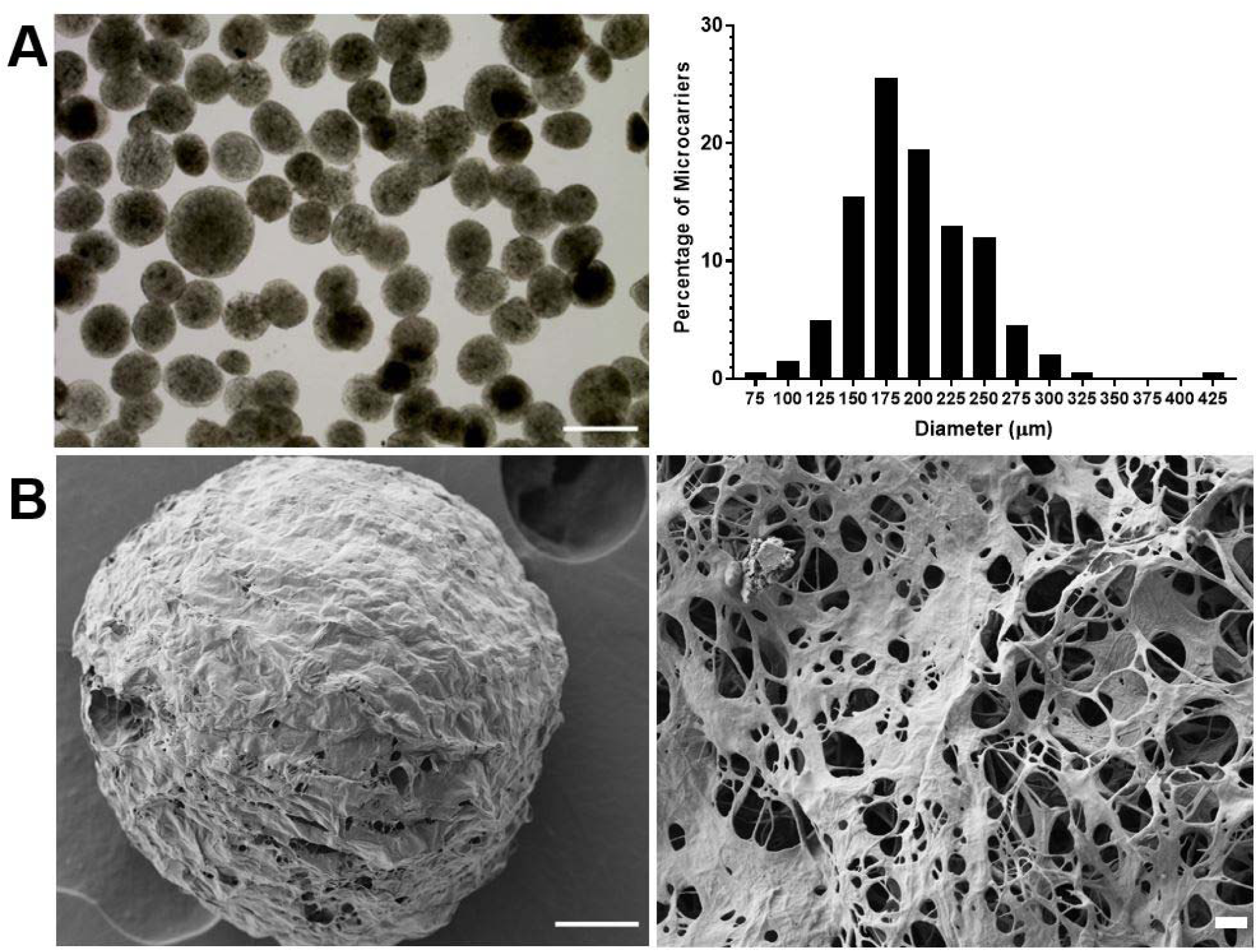
DAT microcarriers fabricated using the microbead generator were stable following rehydration when photo-chemically crosslinked with rose bengal. (A) Representative brightfield image (left) of the DAT microcarriers in PBS (scale bar = 500 µm) and size distribution plot (right) showing the diameter of the rehydrated microcarriers in PBS (n=100 microcarriers/batch, N=2 batches). The mean diameter was 196 ± 47 µm. (B) Representative SEM images showing the microcarrier surface ultrastructure. Scale bar = 50 µm on left, and scale bar = 1 µm on right.

### 3.4 *In vitro* assessment of hASC attachment and growth in spinner flask bioreactors

*In vitro* studies were performed to confirm that the new microcarriers supported hASC attachment and growth within spinner flask bioreactors. An initial trial was performed at a stirring rate of 20 RPM, selected based on our previous studies with our electrosprayed microcarriers [8], [9]. However, the microcarriers were observed to begin aggregating after 7 days of culture under these conditions (Supplementary Fig. S1). In the subsequent trial using the same hASC donor, the stirring rate was increased to 40 RPM, which substantially reduced aggregation and improved cell growth based on analysis of dsDNA content (Supplementary Fig. S1). Additional *in vitro* trials were performed to validate the response across a total of 3 hASC donors. Importantly, the microcarriers were shown to consistently support hASC expansion over 14 days based on LIVE/DEAD® imaging and quantification of dsDNA content (Fig. 5A-D). Analysis of sGAG content showed increased levels in the hASC-seeded microcarriers cultured for 14 days compared to the unseeded controls (Fig. 5E), indicating that the hASCs were producing ECM and remodeling the microcarriers. However, the hydroxyproline content as a measure of total collagen content at day 14 was similar to the unseeded controls (Fig. 5F), consistent with the microcarriers macroscopically maintaining their integrity over the duration of the study.

**Figure 5.**
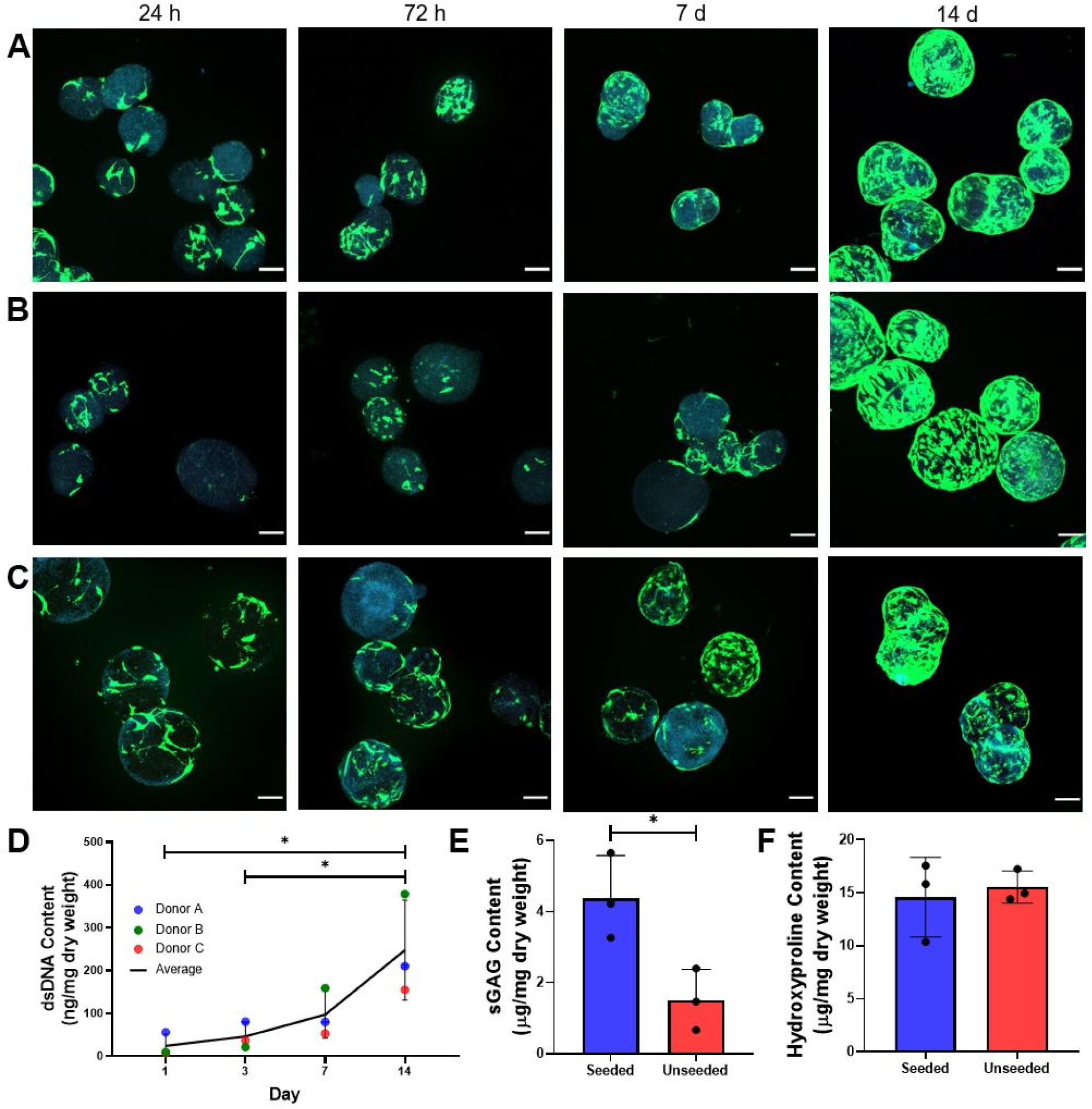
DAT microcarriers supported hASC expansion and sGAG production within spinner flask bioreactors. (A-C) Representative LIVE/DEAD® staining over time of hASCs from 3 different donors seeded on the microcarriers and cultured within spinner flasks at 40 RPM. Green=viable calcein^+^ hASCs, blue=ECM autofluorescence. Scale bars = 100 µm. (D) dsDNA content quantified via the PicoGreen assay showing cell growth over time, with significantly higher dsDNA content at day 14 day compared to days 1 and 3. (E) sGAG content was significantly increased in the seeded samples after 14 days of culture compared to unseeded baseline controls. (F) Hydroxyproline content as a measure of total collagen content was unchanged in the hASC-seeded microcarriers at day 14 compared to baseline unseeded microcarriers. (*p < 0.05, n=3, N=3).

## 4 DISCUSSION

In the current study, we established a new cost-effective method for fabricating ECM-derived microcarriers sourced from decellularized tissues that matched our target diameter of ∼200 μm. Our novel 3D T-junction microbead generator was successful in creating small, monodisperse microbeads with production rates of up to hundreds of beads per second. These beads were stable following photo-crosslinking and rehydration and were successfully used as microcarriers to support the growth of hASCs within spinner flask bioreactors.

Our modified 3D T-junction design has several advantages over previous designs. Its fabrication is cheaper and easier than previous microfluidic T-junction devices. Device fabrication methods previously used required expensive nanofabrication facilities and precise bonding of separate sections [21]. The microfluidic device used in the current study is robust and reusable with a low cost of fabrication, requiring only access to a micro-milling machine. We fabricated the device from acrylic, as it has a high elastic modulus and will not deform under high flow rates, which can induce high pressure within the device. In contrast, polydimethylsiloxane (PDMS), which is commonly used in microfluidic device fabrication [30], is more compliant and may expand noticeably under high pressure, altering the velocity, the drag forces, and the shear stress profiles within the device.

The current design was modified from a 3D T-junction design by Yoon *et al.* [21] to increase shearing rates through the insertion of the dispersed-phase fluid flow into the midstream of the continuous phase. Insertion of the DP needle into the midplane appeared to have a synergistic effect, reducing the cross-sectional area for the CP flow to ∼81% of the original, and thus increasing the mean velocity in the obstructed section by ∼23%. Additionally, the obstructed flow leads to a plug flow profile with higher shearing forces at the tip of the needle. Although T-junctions are prone to clogging, with the current design, the needle can easily be replaced and the entire device opened for cleaning, which was particularly important when working with our viscous α-amylase digested DAT.

Glycolytic digestion with α-amylase was selected as the method of ECM processing as it has been previously shown to have benefits compared to the more common method of proteolytic digestion with pepsin [9], [31]. α-amylase digestion cleaves carbohydrate groups to increase collagen solubility in acetic acid, but better preserves the ECM protein structure and composition, which may be favorable for bioactivity [24]. In a previous study, it was shown that hASC proliferation and adipogenic differentiation were enhanced on coatings fabricated from α-amylase-digested DAT compared to pepsin-digested DAT [32]. However, a challenge from a microfluidics perspective is that α-amylase digestion produces an ECM suspension that is more viscous than pepsin-digested ECM.

Our measurements of the viscosity of the α-amylase digested DAT suspension, at varying DAT concentrations, temperatures, and shear rates, showed that viscosity decreased as a function of shear rate and temperature, ranging from ∼20 mPa⋅s at low temperature and low shear rate to 2 mPa⋅s at high temperature and high shear rate for the 10 mg/mL suspension. Respectively, for the 20 mg/mL concentration, viscosity ranged from 130 mPa⋅s to 7 mPa⋅s, and for the 30 mg/mL concentration, ranged from ∼250 mPa⋅s to 30 mPa⋅s. The dependence of viscosity on shear rate indicates that the DAT suspensions are mildly shear–thinning non-Newtonian fluids. Shear thinning is characteristic of many polymer-based suspensions and reflects the effects of shear on the microstructure of the suspension. Using a DAT concentration as high as practical was important to ensure that the resulting microcarriers were structurally robust and stable in long-term culture. However, increasing the viscosity makes it more challenging to ensure continuous flow when working with small channel sizes. Increasing the temperature and shear rates through increased flow rates are factors that can help alleviate clogging. In our process, the 20 mg/mL DAT suspension was warmed to 25°C to reduce its viscosity. However, the flow rates also have a direct impact on bead size, so this was further explored.

The literature suggests that microparticle size should vary inversely with Q_CP_ with relatively little dependence on Q_DP_ [33]. Our results are in agreement with this, as seen in Fig. 3A. This results in the linear dependence on the flow rate ratio (Q_DP_/Q_CP_) seen in Fig. 3B, although we do observe a dependence on Q_DP_ for the lowest Q_CP_ (Fig. 3A). For Q_CP_ = 1500 µL/min (and all values of Q_DP_), and for Q_CP_ = 500 µL/min and Q_DP_ = 250 µL/min, a bimodal distribution of bead sizes, with secondary beads of ∼50 µm diameter, was produced. Microdroplet formation modelling suggests that the ratio of the bead diameter to CP channel width will approach a limiting value (∼0.3-0.5) at high Q_CP_, with the limit decreasing as the viscosity ratio of the two phases (μ_CP_/μ_DP_) increases [33]. Our viscosity ratio was approximately 2.5, based on a DAT viscosity of ∼40 mPa⋅s at 25°C for the 20 mg/mL suspension and estimating the viscosity of corn oil to be 96 mPa⋅s at ∼4°C [34]. This would suggest we are limited to a larger dimensionless bead size of 0.4-0.45 [33]. Experimentally, however, at our highest value of Q_CP_, the size ratio ranged from 0.24 to 0.34. This suggests our new microfluidic device design allowed us to make beads of comparable size (196 ± 47 μm) to those produced by Yoon *et al*. [21] (175 ± 8 μm), despite our use of a highly viscous 20 mg/mL DAT suspension with lower viscosity ratio than their 2 mg/mL type-I collagen. Previous groups using flow-focusing devices [35], [36] also formed microcarriers similar in diameter with a small size distribution. Therefore, our results are in general agreement with the literature, albeit with some of the microfluidic device design factors perhaps working to our advantage to enable similar sized microbeads despite a larger DP channel size and lower viscosity ratio.

Preferred parameters selected for further characterization were Q_DP_ = 100 µL/min and Q_CP_ = 500 µL/min, as this flow rate ratio (Q_DP_/Q_CP_ = 0.2) was the point at which the monodispersed beads were ∼200 µm in diameter with a high production rate of ∼100 beads/sec. For this flow rate combination, beads were 280 ± 10 µm when measured within the microfluidic device during production. However, once collected, crosslinked, and rehydrated, the DAT microcarriers were found to be smaller than the original microbeads, with an average diameter of 196 ± 47 µm. This reduction in size is potentially attributed to crosslinking [37]. Importantly, the microcarriers were smaller and less variable than the 469 ± 164 μm diameter DAT microcarriers generated by Kornmuller *et al.* through electrospraying [10]. Similar to the electrosprayed microcarriers, those generated by our microfluidic device appeared fairly round and uniform in shape. Homogeneity of microcarrier shape is important, as both size and shape have previously been shown to influence cell yield and viability [38]. Compared to our previous electrosprayed microcarriers, which were not chemically crosslinked [8], those generated via the microfluidic device have a smoother surface ultrastructure, which may be related to continued shearing within the microfluidic device or to crosslinking with rose bengal.

Chemical crosslinking was necessary due to bead aggregation and coalescence after microbead production if the mixture of DAT and oil was disturbed. While glutaraldehyde is commonly used for collagen crosslinking, it has previously been shown to have cytotoxic effects [39]. In an effort to reduce cytotoxicity, we performed photochemical crosslinking using rose bengal [9], [27]. Notably, it was essential to freeze the outflow of the microfluidic device and maintain the mixture between -10°C and -5°C. This temperature allowed the still-frozen beads to maintain their shape, and the rose bengal to penetrate through the liquid corn oil, enabling collagen crosslinking after exposure to visible light.

Photochemical crosslinking has been shown to affect microcarrier stiffness [9], so this was explored. Interestingly, the microcarriers generated using the microfluidic device followed by crosslinking had a similar Young’s modulus compared to those generated by electrospraying (without crosslinking). More specifically, the Young’s modulus of our microcarriers was 0.23 ± 0.05 kPa compared to 0.1 and 0.2 kPa for electrosprayed microcarriers fabricated with 15 mg/mL and 25 mg/mL DAT suspensions [8]. Microcarrier stiffness is an important factor when designing platforms for MSC expansion, as MSCs are sensitive to their mechanical environment [40]. Softer materials may be beneficial, as it has previously been reported that more compliant substrates help to avoid MSC senescence [41]. In addition, the mechanical properties of the ECM are known to affect both MSC proliferation and differentiation and will likely be a key factor in the development of platforms that can maintain an undifferentiated phenotype and augment the pro-regenerative capacity of MSCs during expansion [7], [12], [42].

Initially, cell culture was performed at 20 RPM within the bioreactors. Although the microcarriers supported hASC attachment and growth, substantial microcarrier aggregation was observed. This can be caused by ECM production as well as cell-to-cell interactions [43]. It is important to reduce microcarrier aggregation, as it has been shown to result in decreased cell viability and yields [44]. Strategies to reduce microcarrier aggregation include the addition of fresh microcarriers [43] and increasing the stirring rates [43], [44]. The latter approach was effective with our system. Future studies could further optimize the bioreactor conditions to augment hASC expansion. For example, a previous study assessing hASC growth on polystyrene microcarriers in spinner bioreactors showed that a higher specific growth rate was observed at 49 RPM compared to lower spin speeds of 25 and 43 RPM, with significantly lower growth rates also observed at higher spin speeds of 90 and 120 RPM [45]. Dynamic culture on microcarriers can also modulate the therapeutic potential of MSCs by helping to direct their lineage-specific differentiation [46], as well as modulating the paracrine factors that they produce [47]. This would also be interesting to explore in future work, as would extending the studies to include other MSC sources.

## 5 CONCLUSION

In summary, this study established a novel, cost-effective, and reusable microfluidics-based platform for generating monodisperse DAT-based microcarriers with a diameter of ∼200 µm following stabilization through photo-chemical crosslinking with rose bengal. The micro-milling approach to device fabrication is cheaper and easier than soft lithography techniques. The device produced uniform beads with a high production rate at multiple flow rate combinations. Importantly, these new DAT microcarriers were both smaller and more uniform in size than the DAT microcarriers we previously generated by electrospraying. The microcarriers were shown to support the attachment and growth of hASCs within bioreactor systems, enabling the production of new ECM while maintaining their structural integrity over 14 days. While we established these methods using DAT, which is a readily available human ECM source, they could easily be adapted to generate tissue-specific microcarriers using other decellularized tissue sources, which could be used to support the growth of a wide range of adherent cell types, including MSCs. Given their small size and biological sourcing, these new microcarriers also represent a promising cell delivery platform that could be injected via small gauge needles.

## ACKNOWLEDGMENTS

The authors would like to acknowledge Brian Dalrymple and Frank Van Sas for their machining and technical expertise, and Andrew Roberts for his preliminary microbead formation modelling during prototype design. Dr. Anna Kornmuller and Cherry Liang are also acknowledged for their support in preparing DAT suspensions for the initial device development studies. Drs. Aaron Grant and Damir Matic are acknowledged for their clinical collaborations in providing adipose tissue samples. We would also like to acknowledge Dr. Todd Simpson for technical support and the Western Nanofabrication Facility for access to their infrastructure for SEM imaging. Financial support is acknowledged from the Natural Sciences and Engineering Research Council of Canada Discovery Grant program (RGPIN 06260-2018 and RGPIN-2017-0410) and Research Tools and Instruments program (micro-PIV system); Western University’s Science Undergraduate Summer Research Internship program (FK); the CONNECT! NSERC CREATE Training Program in Soft Connective Tissue Regeneration/Therapy (CJG); and the Ontario Graduate Scholarship (OGS) program (CJG).

## AUTHOR CONTRIBUTIONS

TLP and LEF were responsible for project conceptualization, experimental design, and provided the funding to support this work and contributed to manuscript editing. AA, GB, and JdB contributed to the design and fabrication of the device and the viscosity measurements. FK was responsible for the device implementation, microbead size measurements, devising the cross-linking process, as well as manuscript writing. CJG was responsible for the biochemical characterization of the microcarriers, the cell culture studies, and manuscript writing. TLP supervised the device design, fabrication, and performance characterization. LEF supervised all aspects of the final microcarrier characterization and *in vitro* studies.

## REFERENCES

[1] Z. Weng et al., “Mesenchymal Stem/Stromal Cell Senescence: Hallmarks, Mechanisms, and Combating Strategies,” Stem Cells Transl Med, vol. 11, no. 4, pp. 356–371, Apr. 2022, doi: 10.1093/stcltm/szac004.

[2] M. Lovett, D. Rockwood, A. Baryshyan, and D. L. Kaplan, “Simple Modular Bioreactors for Tissue Engineering: A System for Characterization of Oxygen Gradients, Human Mesenchymal Stem Cell Differentiation, and Prevascularization,” Tissue Eng Part C Methods, vol. 16, no. 6, pp. 1565–1573, Dec. 2010, doi: 10.1089/ten.tec.2010.0241.

[3] J. C. Gerlach et al., “Adipogenesis of Human Adipose-Derived Stem Cells Within Three-Dimensional Hollow Fiber-Based Bioreactors,” Tissue Eng Part C Methods, vol. 18, no. 1, pp. 54– 61, Jan. 2012, doi: 10.1089/ten.tec.2011.0216.

[4] C. Yu, A. Kornmuller, C. Brown, T. Hoare, and L. E. Flynn, “Decellularized adipose tissue microcarriers as a dynamic culture platform for human adipose-derived stem/stromal cell expansion,” Biomaterials, vol. 120, pp. 66–80, Mar. 2017, doi: 10.1016/j.biomaterials.2016.12.017.

[5] T. T. Y. Han, J. T. Walker, A. Grant, G. A. Dekaban, and L. E. Flynn, “Preconditioning Human Adipose-Derived Stromal Cells on Decellularized Adipose Tissue Scaffolds Within a Perfusion Bioreactor Modulates Cell Phenotype and Promotes a Pro-regenerative Host Response,” Front Bioeng Biotechnol, vol. 9, Mar. 2021, doi: 10.3389/fbioe.2021.642465.

[6] A. Mizukami et al., “Proteomic Identification and Time-Course Monitoring of Secreted Proteins During Expansion of Human Mesenchymal Stem/Stromal in Stirred-Tank Bioreactor,” Front Bioeng Biotechnol, vol. 7, Jun. 2019, doi: 10.3389/fbioe.2019.00154.

[7] F. Gattazzo, A. Urciuolo, and P. Bonaldo, “Extracellular matrix: A dynamic microenvironment for stem cell niche,” Biochimica et Biophysica Acta (BBA) - General Subjects, vol. 1840, no. 8, pp. 2506–2519, Aug. 2014, doi: 10.1016/j.bbagen.2014.01.010.

[8] A. Kornmuller and L. E. Flynn, “Development and characterization of matrix□derived microcarriers from decellularized tissues using electrospraying techniques,” J Biomed Mater Res A, vol. 110, no. 3, pp. 559–575, Mar. 2022, doi: 10.1002/jbm.a.37306.

[9] A. E. B. Turner and L. E. Flynn, “Design and Characterization of Tissue-Specific Extracellular Matrix-Derived Microcarriers,” Tissue Eng Part C Methods, vol. 18, no. 3, pp. 186–197, Mar. 2012, doi: 10.1089/ten.tec.2011.0246.

[10] A. Kornmuller, T. T. Cooper, A. Jani, G. A. Lajoie, and L. E. Flynn, “Probing the effects of matrix□derived microcarrier composition on human adipose□derived stromal cells cultured dynamically within spinner flask bioreactors,” J Biomed Mater Res A, vol. 111, no. 3, pp. 415– 434, Mar. 2023, doi: 10.1002/jbm.a.37459.

[11] J. C. Terek, M. O. Hebb, and L. E. Flynn, “Development of Brain-Derived Bioscaffolds for Neural Progenitor Cell Culture,” ACS Pharmacol Transl Sci, vol. 6, no. 2, pp. 320–333, Feb. 2023, doi: 10.1021/acsptsci.2c00232.

[12] C. Yu, A. Kornmuller, C. Brown, T. Hoare, and L. E. Flynn, “Decellularized adipose tissue microcarriers as a dynamic culture platform for human adipose-derived stem/stromal cell expansion,” Biomaterials, vol. 120, pp. 66–80, Mar. 2017, doi: 10.1016/j.biomaterials.2016.12.017.

[13] S. Sart, Y.-J. Schneider, and S. N. Agathos, “Influence of culture parameters on ear mesenchymal stem cells expanded on microcarriers,” J Biotechnol, vol. 150, no. 1, pp. 149–160, Oct. 2010, doi: 10.1016/j.jbiotec.2010.08.003.

[14] T. K.-P. Goh et al., “Microcarrier Culture for Efficient Expansion and Osteogenic Differentiation of Human Fetal Mesenchymal Stem Cells,” Biores Open Access, vol. 2, no. 2, pp. 84–97, Apr. 2013, doi: 10.1089/biores.2013.0001.

[15] W. S. Hu, J. Meier, and D. I. C. Wang, “A mechanistic analysis of the inoculum requirement for the cultivation of mammalian cells on microcarriers,” Biotechnol Bioeng, vol. 95, no. 2, pp. 306–316, Oct. 2006, doi: 10.1002/bit.21157.

[16] P. Silva Couto et al., “Expansion of human mesenchymal stem/stromal cells (hMSCs) in bioreactors using microcarriers: lessons learnt and what the future holds,” Biotechnol Adv, vol. 45, p. 107636, Dec. 2020, doi: 10.1016/j.biotechadv.2020.107636.

[17] A.-C. Tsai, R. Jeske, X. Chen, X. Yuan, and Y. Li, “Influence of Microenvironment on Mesenchymal Stem Cell Therapeutic Potency: From Planar Culture to Microcarriers,” Front Bioeng Biotechnol, vol. 8, Jun. 2020, doi: 10.3389/fbioe.2020.00640.

[18] L. Wu, Z. Guo, and W. Liu, “Surface behaviors of droplet manipulation in microfluidics devices,” Adv Colloid Interface Sci, vol. 308, p. 102770, Oct. 2022, doi: 10.1016/j.cis.2022.102770.

[19] A. B. Theberge et al., “Microdroplets in Microfluidics: An Evolving Platform for Discoveries in Chemistry and Biology,” Angewandte Chemie International Edition, vol. 49, no. 34, pp. 5846–5868, Aug. 2010, doi: 10.1002/anie.200906653.

[20] H. Gu, M. H. G. Duits, and F. Mugele, “Droplets Formation and Merging in Two-Phase Flow Microfluidics,” Int J Mol Sci, vol. 12, no. 4, pp. 2572–2597, Apr. 2011, doi: 10.3390/ijms12042572.

[21] J. Yoon, J. Kim, H. E. Jeong, R. Sudo, M.-J. Park, and S. Chung, “Fabrication of type I collagen microcarrier using a microfluidic 3D T-junction device and its application for the quantitative analysis of cell–ECM interactions,” Biofabrication, vol. 8, no. 3, p. 035014, Aug. 2016, doi: 10.1088/1758-5090/8/3/035014.

[22] P. Morissette Martin, A. Grant, D. W. Hamilton, and L. E. Flynn, “Matrix composition in 3-D collagenous bioscaffolds modulates the survival and angiogenic phenotype of human chronic wound dermal fibroblasts,” Acta Biomater, vol. 83, pp. 199–210, Jan. 2019, doi: 10.1016/j.actbio.2018.10.042.

[23] H. K. Cheung, T. T. Y. Han, D. M. Marecak, J. F. Watkins, B. G. Amsden, and L. E. Flynn, “Composite hydrogel scaffolds incorporating decellularized adipose tissue for soft tissue engineering with adipose-derived stem cells,” Biomaterials, vol. 35, no. 6, pp. 1914–1923, Feb. 2014, doi: 10.1016/j.biomaterials.2013.11.067.

[24] M. Kuljanin, C. F. C. Brown, M. J. Raleigh, G. A. Lajoie, and L. E. Flynn, “Collagenase treatment enhances proteomic coverage of low-abundance proteins in decellularized matrix bioscaffolds,” Biomaterials, vol. 144, pp. 130–143, Nov. 2017, doi: 10.1016/j.biomaterials.2017.08.012.

[25] L. E. Flynn, “The use of decellularized adipose tissue to provide an inductive microenvironment for the adipogenic differentiation of human adipose-derived stem cells,” Biomaterials, vol. 31, no. 17, pp. 4715–4724, Jun. 2010, doi: 10.1016/j.biomaterials.2010.02.046.

[26] D. Cherfan et al., “Collagen Cross-Linking Using Rose Bengal and Green Light to Increase Corneal Stiffness,” Investigative Opthalmology & Visual Science, vol. 54, no. 5, p. 3426, May 2013, doi: 10.1167/iovs.12-11509.

[27] S. Eckes, J. Braun, J. S. Wack, U. Ritz, D. Nickel, and K. Schmitz, “Rose Bengal Crosslinking to Stabilize Collagen Sheets and Generate Modulated Collagen Laminates,” Int J Mol Sci, vol. 21, no. 19, p. 7408, Oct. 2020, doi: 10.3390/ijms21197408.

[28] K. Kim, J. Cheng, Q. Liu, X. Y. Wu, and Y. Sun, “Investigation of mechanical properties of soft hydrogel microcapsules in relation to protein delivery using a MEMS force sensor,” J Biomed Mater Res A, vol. 92A, no. 1, pp. 103–113, Jan. 2010, doi: 10.1002/jbm.a.32338.

[29] A. Shridhar, B. G. Amsden, E. R. Gillies, and L. E. Flynn, “Investigating the Effects of Tissue-Specific Extracellular Matrix on the Adipogenic and Osteogenic Differentiation of Human Adipose-Derived Stromal Cells Within Composite Hydrogel Scaffolds,” Front Bioeng Biotechnol, vol. 7, Dec. 2019, doi: 10.3389/fbioe.2019.00402.

[30] M. P. Wolf, G. B. Salieb-Beugelaar, and P. Hunziker, “PDMS with designer functionalities— Properties, modifications strategies, and applications,” Prog Polym Sci, vol. 83, pp. 97–134, Aug. 2018, doi: 10.1016/j.progpolymsci.2018.06.001.

[31] J. Visser et al., “Crosslinkable Hydrogels Derived from Cartilage, Meniscus, and Tendon Tissue,” Tissue Eng Part A, vol. 21, no. 7–8, pp. 1195–1206, Apr. 2015, doi: 10.1089/ten.tea.2014.0362.

[32] A. Shridhar, A. Y. L. Lam, Y. Sun, C. A. Simmons, E. R. Gillies, and L. E. Flynn, “Culture on Tissue□Specific Coatings Derived from α□Amylase□Digested Decellularized Adipose Tissue Enhances the Proliferation and Adipogenic Differentiation of Human Adipose□Derived Stromal Cells,” Biotechnol J, vol. 15, no. 3, Mar. 2020, doi: 10.1002/biot.201900118.

[33] J. Husny and J. J. Cooper-White, “The effect of elasticity on drop creation in T-shaped microchannels,” J Nonnewton Fluid Mech, vol. 137, no. 1–3, pp. 121–136, Aug. 2006, doi: 10.1016/j.jnnfm.2006.03.007.

[34] S. N. Sahasrabudhe, V. Rodriguez-Martinez, Meghan. O’Meara, and B. E. Farkas, “Density, viscosity, and surface tension of five vegetable oils at elevated temperatures: Measurement and modeling,” Int J Food Prop, pp. 1–17, Dec. 2017, doi: 10.1080/10942912.2017.1360905.

[35] J. S. Lee et al., “Tissue Beads: Tissue□Specific Extracellular Matrix Microbeads to Potentiate Reprogrammed Cell□Based Therapy,” Adv Funct Mater, vol. 29, no. 31, Aug. 2019, doi: 10.1002/adfm.201807803.

[36] L. Yu et al., “Core-shell hydrogel beads with extracellular matrix for tumor spheroid formation,” Biomicrofluidics, vol. 9, no. 2, Mar. 2015, doi: 10.1063/1.4918754.

[37] B. Sridhar, A. Srinatha, and M. Khan, “Development and evaluation of pH-dependent micro beads for colon targeting,” Indian J Pharm Sci, vol. 72, no. 1, p. 18, 2010, doi: 10.4103/0250-474X.62230.

[38] A. K.-L. Chen, X. Chen, A. B. H. Choo, S. Reuveny, and S. K. W. Oh, “Critical microcarrier properties affecting the expansion of undifferentiated human embryonic stem cells,” Stem Cell Res, vol. 7, no. 2, pp. 97–111, Sep. 2011, doi: 10.1016/j.scr.2011.04.007.

[39] M. E. Nimni, D. Cheung, B. Strates, M. Kodama, and K. Sheikh, “Chemically modified collagen: A natural biomaterial for tissue replacement,” J Biomed Mater Res, vol. 21, no. 6, pp. 741–771, Jun. 1987, doi: 10.1002/jbm.820210606.

[40] H. I. Gungordu, M. Bao, S. Helvert, J. A. Jansen, S. C. G. Leeuwenburgh, and X. F. Walboomers, “Effect of mechanical loading and substrate elasticity on the osteogenic and adipogenic differentiation of mesenchymal stem cells,” J Tissue Eng Regen Med, vol. 13, no. 12, pp. 2279–2290, Dec. 2019, doi: 10.1002/term.2956.

[41] R. C. H. Gresham et al., “Compliant substrates mitigate the senescence associated phenotype of stress induced <scp>mesenchymal stromal cells</scp>,” J Biomed Mater Res A, vol. 112, no. 5, pp. 770–780, May 2024, doi: 10.1002/jbm.a.37657.

[42] J. Lee, A. A. Abdeen, A. S. Kim, and K. A. Kilian, “Influence of Biophysical Parameters on Maintaining the Mesenchymal Stem Cell Phenotype,” ACS Biomater Sci Eng, vol. 1, no. 4, pp. 218–226, Apr. 2015, doi: 10.1021/ab500003s.

[43] C. Ferrari, F. Balandras, E. Guedon, E. Olmos, I. Chevalot, and A. Marc, “Limiting cell aggregation during mesenchymal stem cell expansion on microcarriers,” Biotechnol Prog, vol. 28, no. 3, pp. 780–787, May 2012, doi: 10.1002/btpr.1527.

[44] J. Lembong et al., “Bioreactor Parameters for Microcarrier-Based Human MSC Expansion under Xeno-Free Conditions in a Vertical-Wheel System,” Bioengineering, vol. 7, no. 3, p. 73, Jul. 2020, doi: 10.3390/bioengineering7030073.

[45] V. Jossen et al., “Theoretical and Practical Issues That Are Relevant When Scaling Up hMSC Microcarrier Production Processes,” Stem Cells Int, vol. 2016, no. 1, Jan. 2016, doi: 10.1155/2016/4760414.

[46] J. E. Frith, B. Thomson, and P. G. Genever, “Dynamic Three-Dimensional Culture Methods Enhance Mesenchymal Stem Cell Properties and Increase Therapeutic Potential,” Tissue Eng Part C Methods, vol. 16, no. 4, pp. 735–749, Aug. 2010, doi: 10.1089/ten.tec.2009.0432.

[47] R. Jeske et al., “Agitation in a microcarrier-based spinner flask bioreactor modulates homeostasis of human mesenchymal stem cells,” Biochem Eng J, vol. 168, p. 107947, Apr. 2021, doi: 10.1016/j.bej.2021.107947.

