## Supplementary Fig. S1 for "A simple, cost-effective microfluidic device using a 3D cross-flow T-junction for producing decellularized extracellular matrix-derived microcarriers"

**SUPPLEMENTAL MATERIAL**

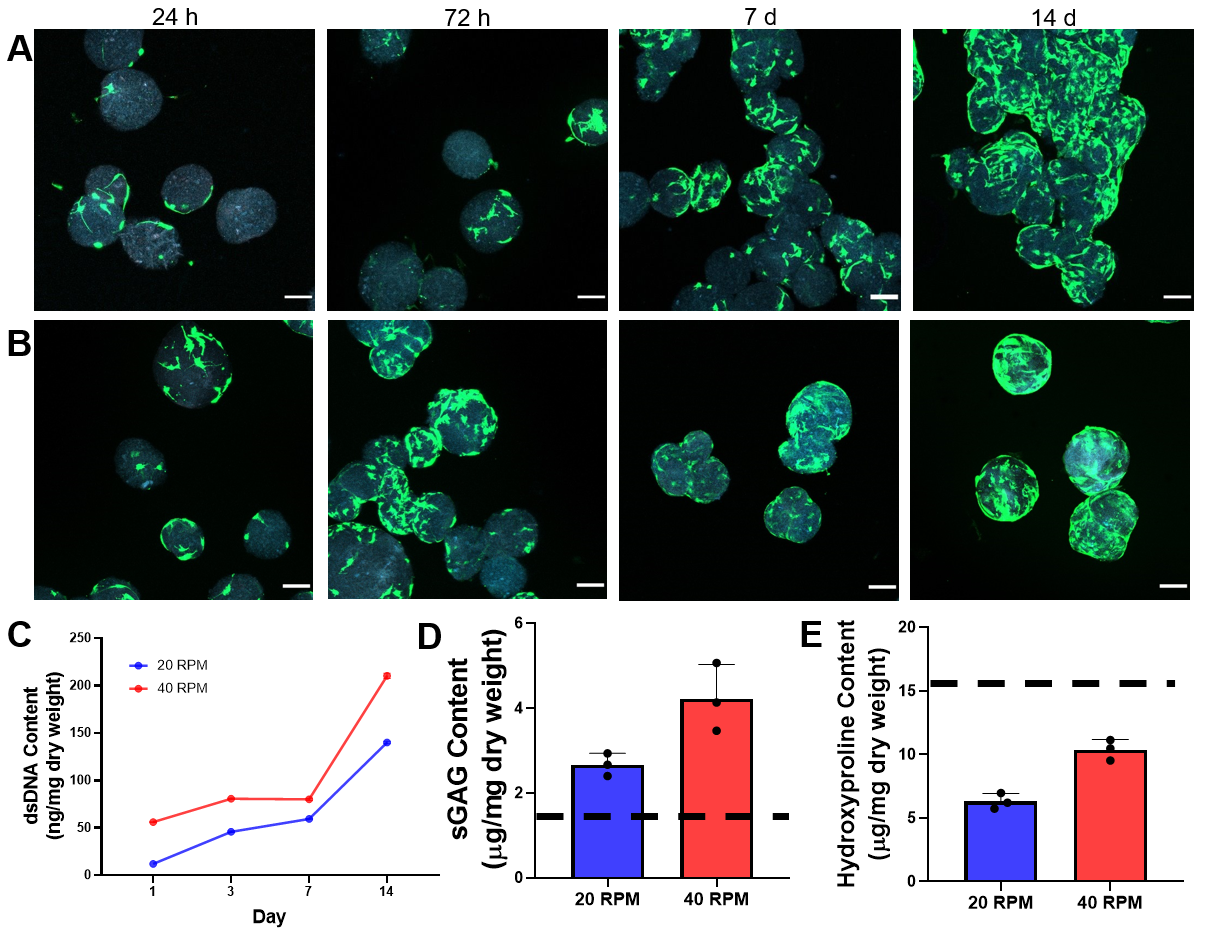

**Figure S1. DAT microcarriers supported hASC expansion over 14 days within spinner-flask bioreactors at 20 RPM and 40 RPM, with microcarrier aggregation observed at 20 RPM.** (A) Representative LIVE/DEAD® staining of microcarriers cultured within spinner flasks at 20 RPM and (B) 40 RPM, showing the formation of large aggregates after 14 days of culture at 20 RPM. Green=viable calcein^+^ hASCs, blue=ECM autofluorescence. Scale bars = 100 µm. (C) dsDNA content quantified via the PicoGreen assay, showing increased cellularity over 14 days. (D) sGAG content at 20 RPM and 40 RPM compared to unseeded controls (dashed line). (E) Hydroxyproline content as a measure of collagen content at 20 RPM and 40 RPM compared to unseeded controls (dashed line). (n=3, N=1).

**TABLE S1: Fit parameters for Cross model (Eqn. 1) fit to viscosity data shown in Fig. 2.** Values represent mean and standard deviation based on the 95% confidence intervals.

| Sample | | $\boldsymbol{\mu}_{\boldsymbol{0}}$ (x 10^3^ mPa·s) | $\boldsymbol{\mu}_{\boldsymbol{\infty}}$(mPa·s) | $\boldsymbol{k}$ | $\boldsymbol{n}$ |
| --- | --- | --- | --- | --- | --- |
| 10 mg/mL | 4°C | 20 ± 10 | 6 ± 3 | (5 ± 3) x 10^4^ | 0.8 ± 0.4 |
|  | 15°C | 60 ± 30 | 5 ± 2 | (6 ± 3) x 10^4^ | 0.9 ± 0.5 |
|  | 22°C | 2 ± 1 | 4 ± 2 | (2 ± 1) x 10^5^ | 0.6 ± 0.3 |
|  | 37°C | 9 ± 4 | 2 ± 2 | (4 ± 2) x 10^4^ | 0.9 ± 0.4 |
| 20 mg/mL | 4°C | 2 ± 1 | 13 ± 6 | (2 ± 1) x 10^3^ | 0.4 ± 0.2 |
|  | 15°C | 0.1 ± 0.1 | 13 ± 7 | 4 ± 2 | 0.6 ± 0.3 |
|  | 22°C | 2 ± 1 | 11 ± 5 | (4 ± 2) x 10^4^ | 0.4 ± 0.2 |
|  | 37°C | 4 ± 2 | 6 ± 3 | (9 ± 4) x 10^4^ | 0.5 ± 0.3 |
| 30 mg/mL | 4°C | 0.3 ± 0.2 | 0 | 0.4 ± 0.2 | 0.4 ± 0.2 |
|  | 15°C | 0.3 ± 0.1 | 0 | 0.2 ± 0.9 | 0.5 ± 0.2 |
|  | 22°C | 0.3 ± 0.1 | 0 | 0.7 ± 0.4 | 0.4 ± 0.2 |
|  | 37°C | 0.2 ± 0.1 | 0 | 0.2 ± 0.1 | 0.4 ± 0.2 |
